# Gene Expression and Tracer-Based Metabolic Flux Analysis Reveals Tissue-Specific Metabolic Scaling *in vitro, ex vivo*, and *in vivo*

**DOI:** 10.1101/2022.03.02.482685

**Authors:** Ngozi D. Akingbesote, Brooks P. Leitner, Daniel G. Jovin, Reina Desrouleaux, Wanling Zhu, Zongyu Li, Michael N. Pollak, Rachel J. Perry

**Affiliations:** Departments of Cellular & Molecular Physiology, Yale School of Medicine, New Haven, CT 06510; Departments of Internal Medicine – Endocrinology, Yale School of Medicine, New Haven, CT 06510; Departments of Comparative Medicine, Yale School of Medicine, New Haven, CT 06510; Lady Davis Institute for Medical Research, Jewish General Hospital, Montreal, QC, Canada; Department of Oncology, McGill University, Montreal, QC, Canada

**Author notes:** Equal contributions (Ngozi D. Akingbesote, Brooks P. Leitner).

## Abstract

Metabolic scaling, the inverse correlation of metabolic rates to body mass, has been appreciated for more than 80 years. Studies of metabolic scaling have almost exclusively been restricted to mathematical modeling of oxygen consumption. The possibility that other metabolic processes scale with body size has not been studied. To address this gap in knowledge, we employed a systems approach spanning from transcriptomics to *in vitro* and *in vivo* tracer-based flux. Gene expression in livers of five species spanning a 30,000-fold range in mass revealed differential expression of genes related to cytosolic and mitochondrial metabolic processes, in addition to detoxication of oxidative damage. This suggests that transcriptional scaling of damage control mechanisms accommodates increased oxidative metabolism in smaller species. To determine whether flux through key implicated metabolic pathways scaled, we applied stable isotope tracer methodology to study multiple cellular compartments, tissues, and species. Comparing mice and rats, we demonstrate that while scaling of metabolic fluxes is not observed in the cell-autonomous setting, it is present in liver slices and *in vivo*. Together, these data reveal that metabolic scaling extends beyond oxygen consumption to numerous other metabolic pathways, and is likely regulated at the level of gene expression and substrate supply.

## Introduction

In 1932, Max Kleiber published a seminal study (Kleiber, 1932), integrating prior reports demonstrating a phenomenon that came to be termed “Kleiber’s law,” or the principle of metabolic scaling. Metabolic scaling refers to the phenomenon that the metabolic rates of many animals, if not all, scale inversely to three-quarters of their body mass (West et al., 1997). In simpler terms, there is a reduction in metabolic rate as body size increases. For example, an elephant is 25 million times larger than a fruit fly, yet its energy expenditure is only 20 thousand times higher; thus, from fruit fly to elephant, the metabolic rate per gram of body weight scales down 1,250 times. Many hypotheses have been proposed to explain the source of this phenomenon. These hypotheses can be divided into two categories: the first is based on the internal determinants of metabolic rate – based on the animal’s internal physical constraints such as the fractal transportation that determines nutrient and oxygen supply (Darveau et al., 2002) and utilization (Darveau et al., 2005) rates across the animal’s body surfaces and internal organs (Savage et al., 2007; Thommen et al., n.d.; West et al., 1997), and the second emphasizes the external determinants of metabolic rate – based on ecological restraints on metabolism, such as the optimization of reproductive success based on size-specific rates of survival, which increases markedly in larger animals as compared to smaller (Thommen et al., n.d.). However, these hypotheses have been derived almost exclusively from observations of caloric intake and oxygen consumption, with gene expression, and substrate fluxes including lipolysis, gluconeogenesis, and substrate contributions to mitochondrial oxidation almost entirely unexplored.

The theory of hierarchical regulation, that is that gene expression initiates the cascade that allows for flux of metabolic pathways,(Rossell et al., 2005; Suarez and Moyes, 2012) provides a systems framework to understand scaling. Due to the relatively recent advances in high throughput mRNA sequencing and bioinformatics tools that allow for intra-specific data preprocessing (Bray et al., 2016; Conesa et al., 2016; Ritchie et al., 2015), we searched for a set of genes whose expression follows the pattern of metabolic scaling in the liver, the metabolic hub of mammals. Since at basal levels, gene expression of metabolic genes closely approximates protein levels (Schwanhäusser et al., 2011), we hypothesized that genes related to the regulation of oxygen consumption and substrate metabolism may scale, and thus provide a mechanistic transcriptional basis for metabolic scaling across five species: mice (*mus musculus*), rats (*rattus norvegicus*), monkeys (*macaca mulatta*), humans (*homo sapiens*), and cattle (*bos taurus*), species with a 30,000-fold range of average body weight in adults (from 30 g in mice, to 900 kg in cattle). Numerous metabolic genes related to glycolysis, gluconeogenesis, fatty acid metabolism, oxygen consumption, electron transport, and redox function, and detoxification of oxidative damage, followed the pattern of metabolic scaling, and informed our isotope-tracer based *in vitro and in vivo* metabolic flux studies.

To determine if gene expression would correlate with metabolic flux, and therefore a mechanistic read out of metabolic enzyme function, the phenomenon of metabolic scaling our lab generated a comprehensive assessment of liver metabolism *in vivo* and *in vitro* using modified Positional Isotopomer NMR Tracer Analysis (PINTA) (Perry et al., 2017b) and stable isotope-derived turnover (Perry et al., 2015) methods to measure mitochondrial oxidation of glucose and fatty acids, gluconeogenesis from pyruvate and glycerol, and white adipose tissue lipolysis in rats and mice at multiple nodes of complexity: *in vitro* in hepatocytes; *ex vivo* in liver slices comprised of hepatocytes plus the surrounding stellate, Kupffer, and sinusoidal endothelial cells; and *in vivo* in conscious mice. Our analysis shows that rats exhibit lower metabolic rates when compared to mice *in* and *ex vivo*; however, no significant differences were observed when we isolated hepatocytes and cultured them *in vitro* under identical conditions.

Taken together, this study demonstrates systems regulation of metabolic scaling: gene expression in livers showed that scaling occurs to regulate oxygen consumption and substrate supply, isotope-based tracer studies in mice and rats demonstrated the mechanistic function of these enzymes *in vivo* which was only apparent in the living organism rather than plated cells. This study provides unique insight into the metabolic scaling at the level of gene expression and metabolic enzyme function.

## Results

### Genes that scale within the liver are predominantly metabolic genes

Considering that mRNA expression correlates well with protein expression under basal conditions, especially for metabolic genes (Schwanhäusser et al., 2011), we used mRNA expression as a proxy for the relative abundance of metabolic enzymes. We examined gene expression in livers from mice (*mus musculus*), rats (*rattus norvegicus*), monkeys (*macaca mulatta*), humans (*homo sapiens*), and cattle (*bos taurus*). After normalizing for differences in transcript length and abundance across species, we filtered out genes that followed the pattern of metabolic scaling. Genes that followed the pattern of mouse > rat > monkey > human > cow were predominantly related to metabolic pathways, including pyruvate metabolism, amino acid metabolism, and glucose metabolism (**Figure 1A)**. Genes from this scaling list were further filtered down to genes involved in amino acid, carbohydrate, energy, lipid, vitamin, and TCA cycle metabolism, and demonstrated a range of degrees of scaling, with only TCA cycle genes clustering together (**Figure 1B)**.

**Figure 1.**
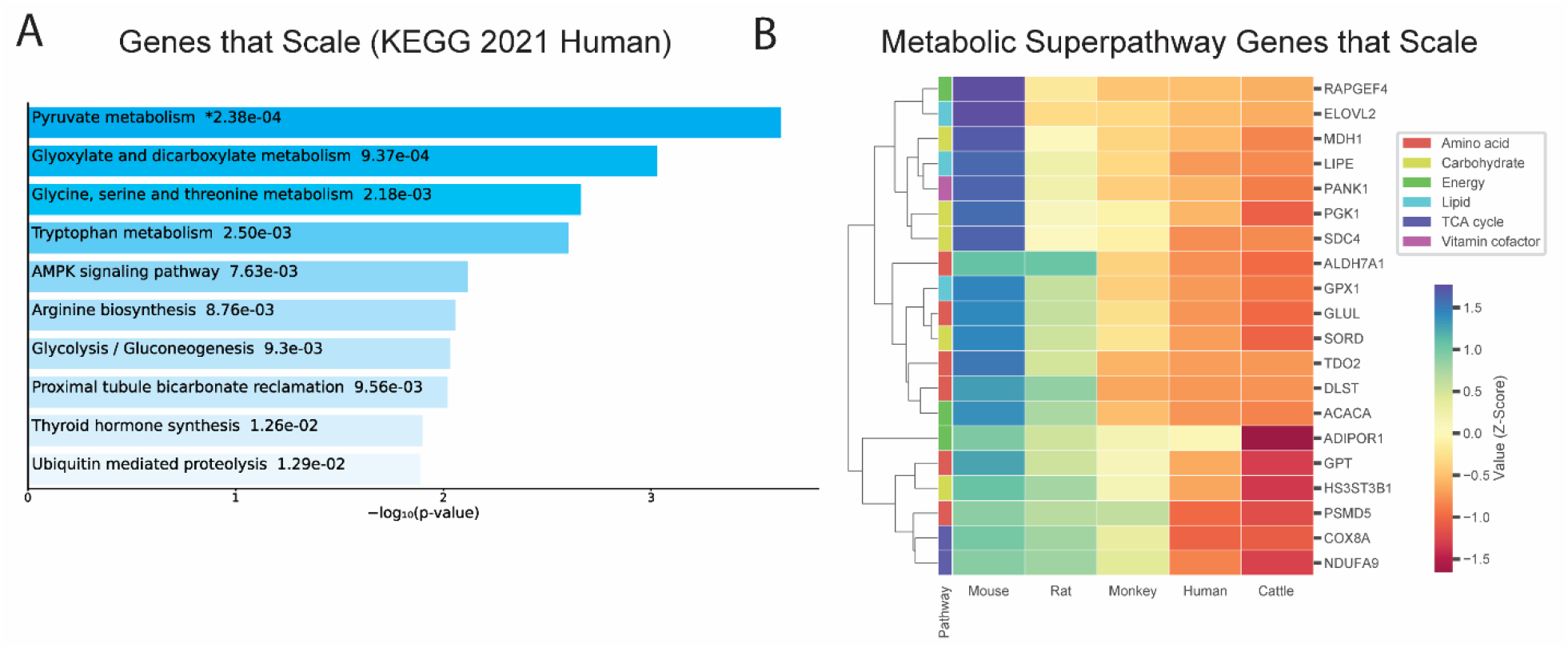
Genes that follow pattern of allometric scaling are most strongly related to metabolism. (A) KEGG Pathway enrichment of all genes that follow pattern of allometric scaling, and (B) clustering heatmap of scaled genes that belong to one of six Reactome metabolic superpathways.

### Metabolic genes that scale in the liver correspond to metabolite detoxification, intertissue metabolism, substrate metabolism, electron transport chain, and NAD metabolism

In order to further understand the functional aspects of the metabolic genes that scale, the gene list from Figure 1B was categorized into several functional categories, converging on optimizing energy provision, oxidative metabolism, and damage control from oxidative stress and ammonia (**Figure 2)**. Further, to understand whether or not certain genes that scale were energy consumers or suppliers themselves, they were further classified into requirements of enzyme action. Eleven of sixteen critical metabolic enzymes that scaled required molecular oxygen, NAD+/NADH, or ATP/ADP for function, possibly indicating exquisite regulation of energy-consuming processes on the individual gene level. Genes involved in detoxication of lipid peroxidation-derived aldehydes (ALDH7A1), hydrogen peroxide (GPX1), and ammonia (GLUL) suggest scaling of damage control mechanisms that are associated with increased oxidative metabolism across species (**Figure 2A)**. Scaling of genes that are associated with interorgan crosstalk provide evidence for mechanisms of scaling *in vivo*, that would not be apparent in plated cells: GPT1, which is involved in recycling skeletal muscle-derived alanine back to glucose through the glucose-alanine cycle (Felig and Wahren, 1971; Petersen et al., 2019), and the adiponectin receptor (ADIPOR1), that binds an adipose tissue-derived hormone, adiponectin, that regulates gluconeogenesis and fatty acid oxidation (Lin and Accili, 2011) **(Figure 2B)**. Genes involved in fatty acid metabolism included the rate limiting steps of the synthesis of CoA (PANK1), of de novo fatty acid synthesis (ACACA), and of fatty acid elongation (ELOVL2), in addition to oxidation of diacylglycerols (LIPE) (**Figure 2C**). NAD and ATP-dependent genes involved in glycolysis (PDK1), fructose/glucose metabolism (SORD1), and DLST of the TCA cycle also scaled (**Figure 2D-E**). Multiple mechanisms are suggested to example oxygen consumption at the level of NAD provision for (via the malate/aspartate shuttle (Williamson et al., 1967) catalyzed by MDH1, and tryptophan catabolism to NAD initiated by TDO2), and function of the electron transport chain (subunits of complex I, NDUFA9, and complex IV, COX8A, which catalyzes oxygen accepting the final electrons of the electron transport chain) **(Figure 2F-G)** (Martínez-Reyes and Chandel, 2020).

**Figure 2.**
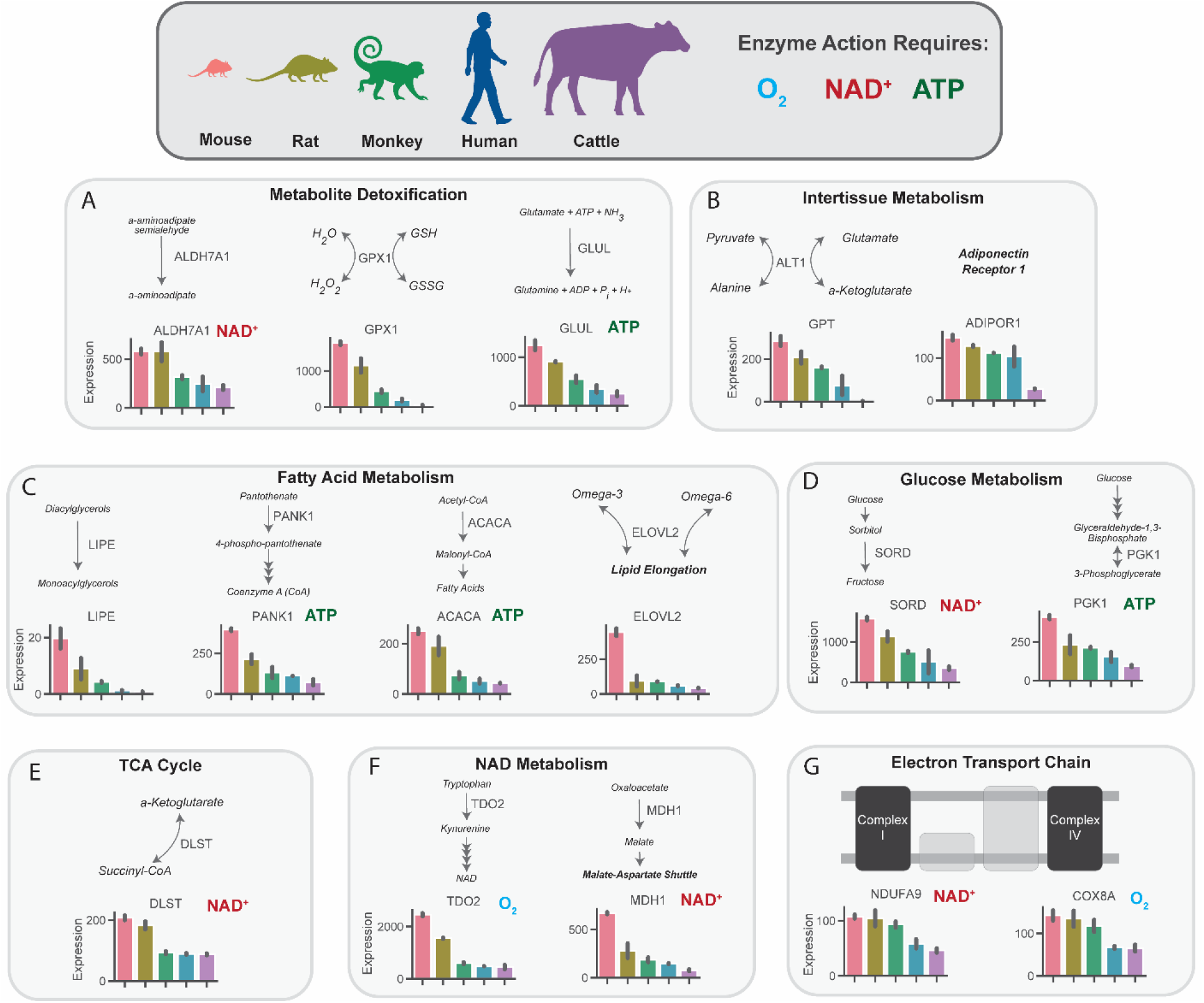
Metabolic genes that scale implicate key pathways in substrate and nucleotide supply, glucose and fatty acid flux, oxygen consumption, and detoxification pathways. mRNA expression of key regulatory genes related to metabolite detoxication (A), intertissue metabolism (B), fatty acid metabolism (C), glucose metabolism (D), tricarboxylic acid (TCA) cycle, NAD metabolism (F), and the electron transport chain (G) in mice, rats, monkeys, humans, and cattle. All genes met an adjusted p-value threshold of 0.01 using a one-way ANOVA with the Bonferroni correction for multiple comparisons.

### Metabolic rates are not significantly different *in vitro* in plated hepatocytes from mice as compared to rats

Considering data reporting marked increases in systemic oxygen consumption in smaller as compared to larger animals, we first asked whether these differences were characteristic of hepatocytes *per se* or whether *in vivo* or hepatocyte-extrinsic signals are required. Consistent with the latter possibility, when we incubated plated hepatocytes in [3-^13^C] lactate and used PINTA to assess a broad spectrum of cytosolic and mitochondrial fluxes, we observed no significant differences in any of the fluxes measured in plated hepatocytes: glucose production, V_PC_, V_CS_, the contribution of glucose or fatty acids to the tricarboxylic acid (TCA) cycle, or lipolysis (**Figure 3A-H, Figure S1**). Similarly, a mitochondrial stress test in plated hepatocytes revealed no difference in any parameter: neither basal mitochondrial and non-mitochondrial respiration, ATP production, maximal (uncoupled) respiration, spare respiratory capacity, nor proton leak were different between plated hepatocytes from mice and rats (**Figure 3I**). Previous studies have demonstrated scaling at the *in vitro* level in cell suspensions only when analyzed immediately after hepatocyte isolation (Porter and Brand, 1995), and have suggested that the phenomenon of scaling gradually disappears around 24-hours post removal (Brown et al., 2007), similar to the conditions in which we performed these studies, though the majority of *in vitro* studies have also demonstrated an absence of scaling (Glazier, 2015). We extend these results by adding gluconeogenic and lipolytic fluxes in hepatocytes, glucose production in liver slices, and comprehensive flux analysis *in vivo*.

**Figure 3.**
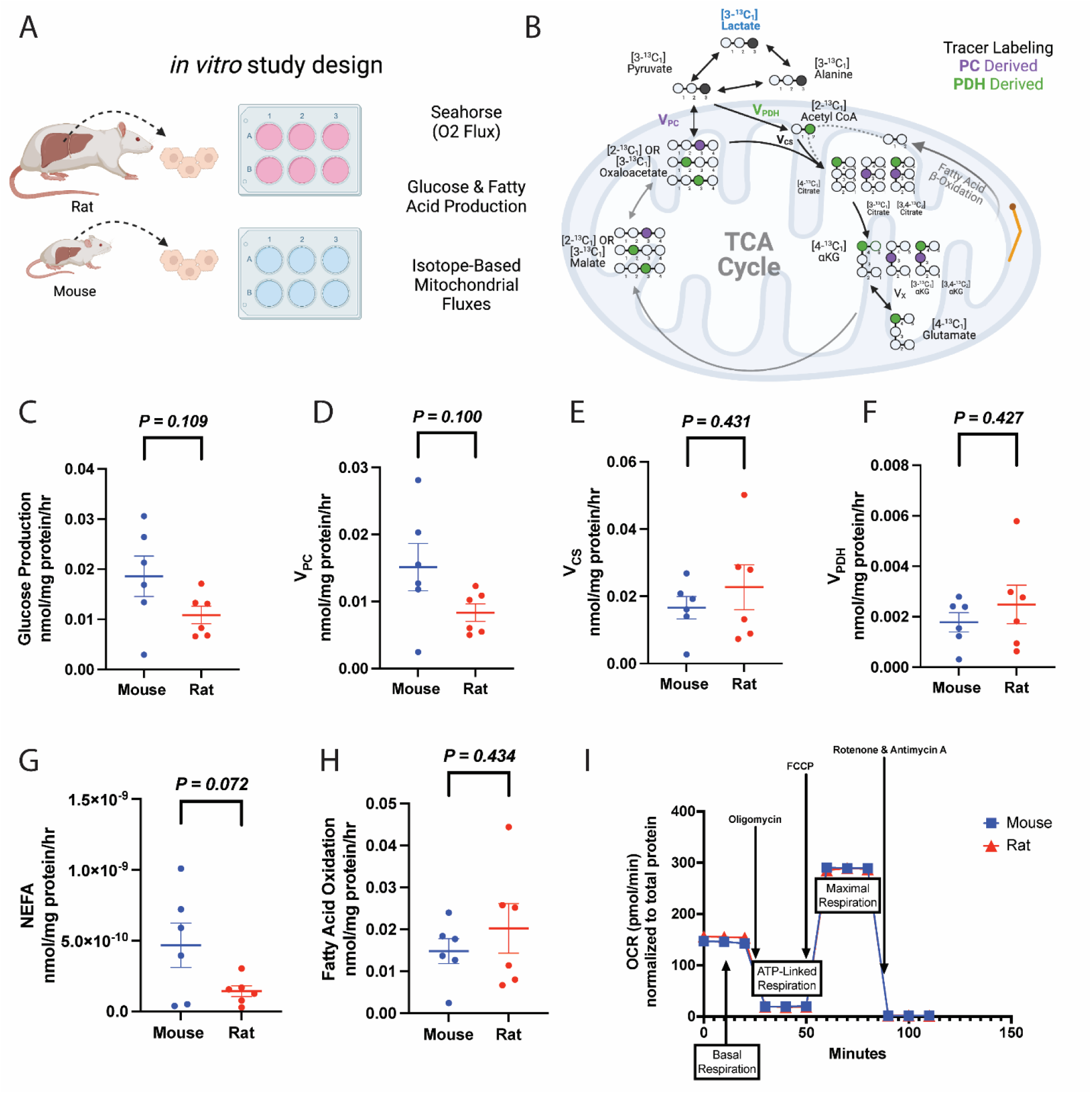
Metabolic fluxes do not scale *in vitro* in hepatocytes. (A) Study design. This figure was made using Biorender.com. (B) Tracer labeling strategy. (C) Glucose production. (D) Gluconeogenesis from pyruvate (pyruvate carboxylase flux, V_PC_). (E) Citrate synthase flux (V_CS_), i.e. mitochondrial oxidation. (F) Pyruvate dehydrogenase flux (V_PDH_), i.e. the contribution of glucose via glycolysis to total mitochondrial oxidation. (G) Non-esterified fatty acid (NEFA) production. (H) The contribution of fatty acid oxidation to citrate synthase flux. (I) Oxygen consumption rate (OCR) during a mitochondrial stress test. In all panels, groups were compared using the 2-tailed unpaired Student’s t-test. No significant differences were observed.

### Glucose production increases *ex vivo* in liver slices in mice relative to rats

Next, considering that hepatocytes comprise approximately 70-80% of liver mass and that their isolation and culture *in vitro* does not replicate *in vivo* conditions (Krebs, 1950), we asked whether glucose production would scale between mice and rats. Indeed, we found that liver glucose production was three-fold greater in mouse liver slices as compared to rat (**Figure 4A- B**), demonstrating metabolic scaling of gluconeogenesis, and suggesting that hepatocyte- extrinsic signals are involved in liver metabolic scaling.

**Figure 4.**
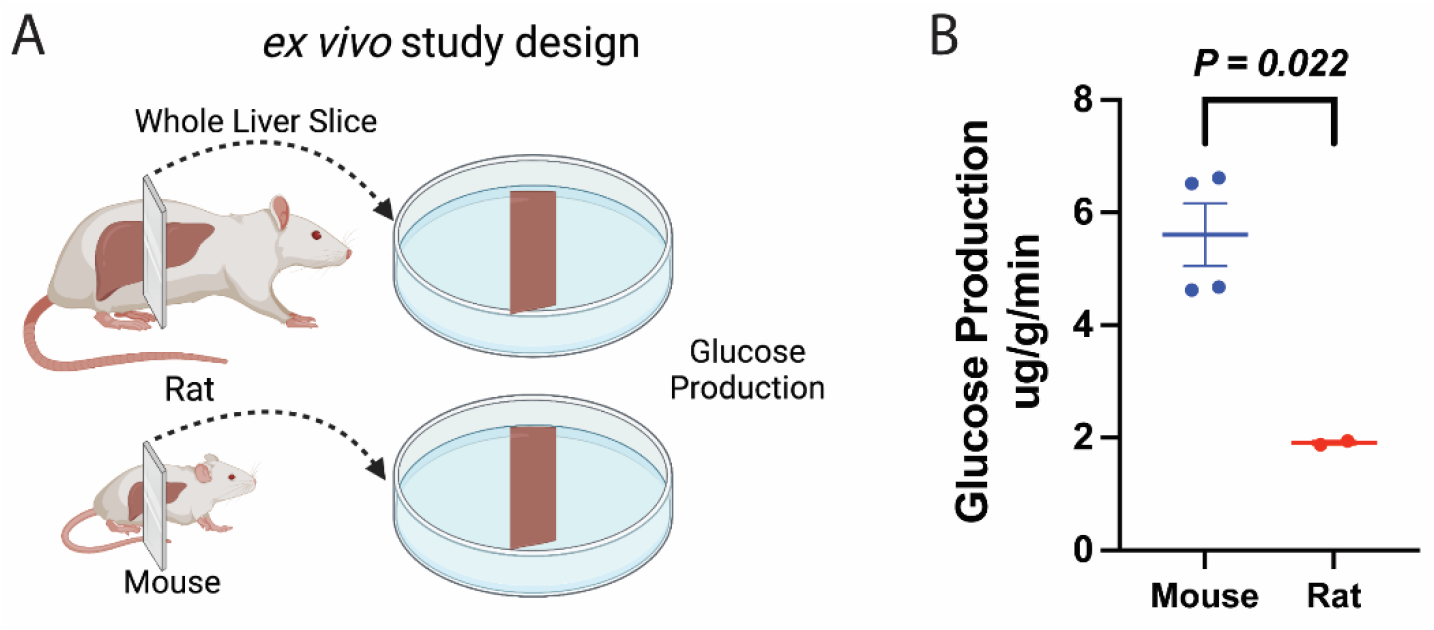
Glucose production scales *ex vivo* in liver slices. (A) Study design. This figure was made using Biorender.com. (B) Glucose production. Groups were compared by the 2-tailed unpaired Student’s t-test.

### Metabolic rates in multiple tissue types are higher *in vivo* in mice relative to rats

We utilized multimodal stable isotope metabolic flux analysis to examine a comprehensive panel of metabolic flux rates in mice and rats (**Figure 5A**). Using PINTA, we found that both endogenous glucose production and gluconeogenesis from pyruvate (pyruvate carboxylase flux, V_PC_) per gram liver were more than twofold higher in mice than rats (**Figure 5B-C**), although the fractional contribution of pyruvate to gluconeogenesis did not differ between mice and rats.

**Figure 5.**
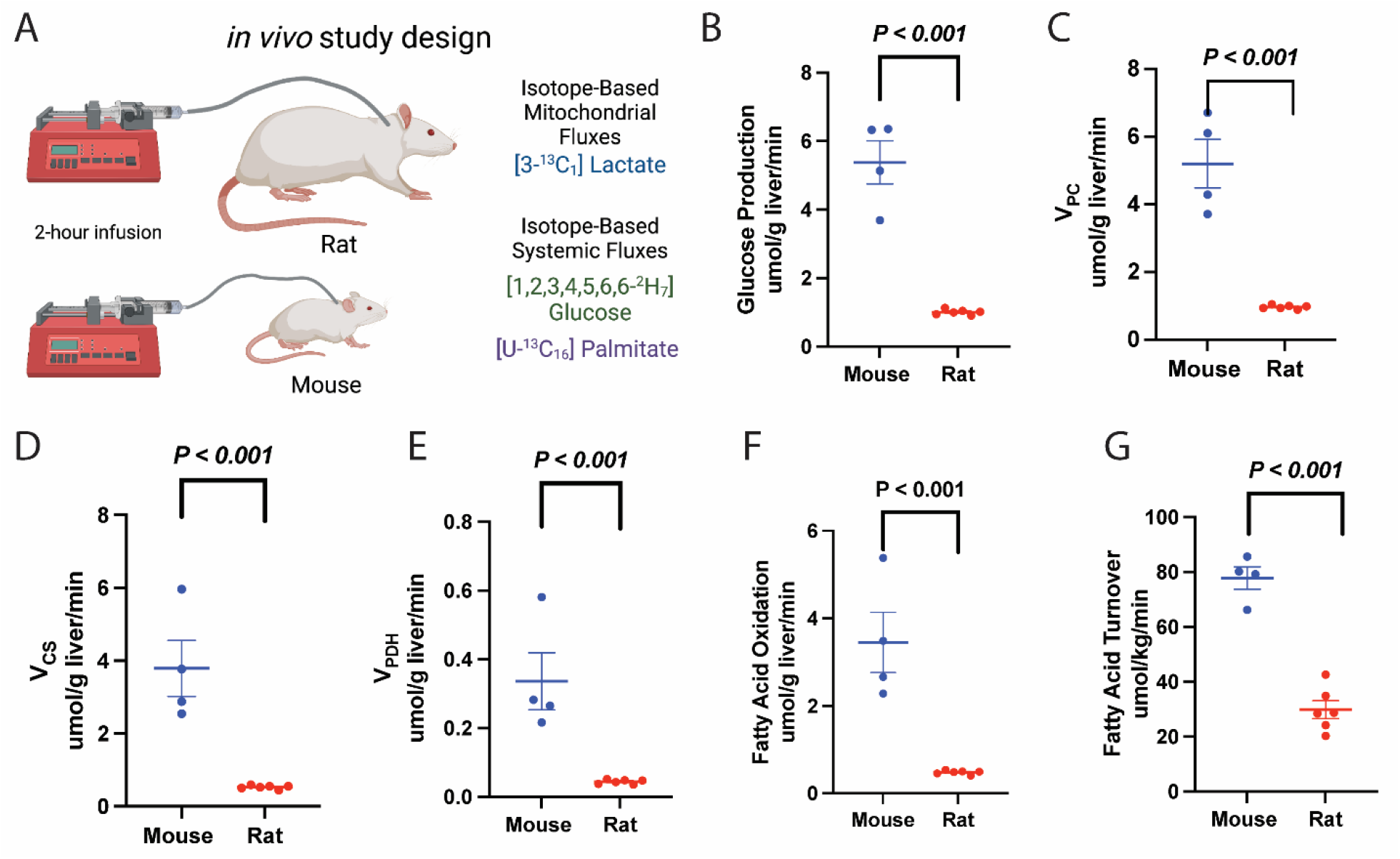
A comprehensive analysis of systemic metabolic fluxes reveals *in vivo* scaling in mice versus rats. (A) Study design. (B) Endogenous glucose production. (C) Gluconeogenesis from pyruvate (V_PC_). (D) V_CS_, i.e. mitochondrial oxidation. (E) V_PDH_, i.e. the contribution of glucose via glycolysis to total mitochondrial oxidation. (F) Palmitate (fatty acid) turnover. (G) The contribution of fatty acid oxidation to citrate synthase flux. In all panels, groups were compared using the 2-tailed unpaired Student’s t-test.

Mitochondrial oxidation scaled similarly, increasing threefold in mice as compared to rats studied under the same conditions, due to increases in both glucose oxidation (pyruvate dehydrogenase flux, V_PDH_) and fatty acid oxidation (**Figure 5D-F**), without any difference in the ratio of pyruvate carboxylase anaplerosis to citrate synthase flux (V_PC_/V_CS_) or the fraction of V_CS_ flux fueled by glucose through PDH (**Figure S2**). This conservation of metabolic scaling applied not only to hepatocyte metabolism but also to adipose tissue metabolism: whole-body fatty acid turnover, reflecting lipolysis, was 2.5-fold higher in mice than rats (**Figure 5G**), demonstrating coordination of systemic metabolic scaling in mice and rats. These data emphasize the inadequacy of common *in vitro* methods as a readout of *in vivo* metabolism: whereas *in vivo* mitochondrial oxidation (TCA cycle flux) was threefold higher in mice than in rats, *in vitro* measurements of oxygen consumption throughout a mitochondrial stress test, TCA cycle flux, and glucose production were not different between the species (**Figure 3**).

### Machine learning defined species-specific clusters based on *in vivo* metabolic fluxes but not *in vitro* fluxes

A clustering dendrogram was applied to our *in vitro* metabolic scaling data and showed no distinct clustering between species (**Figure 6A-B)**. However, the *in vivo* metabolic flux data led to distinct clustering of rats and mice (**Figure 6C)**, providing a machine learning-based objective analysis of *in vitro* versus *in vivo* metabolic flux.

**Figure 6.**
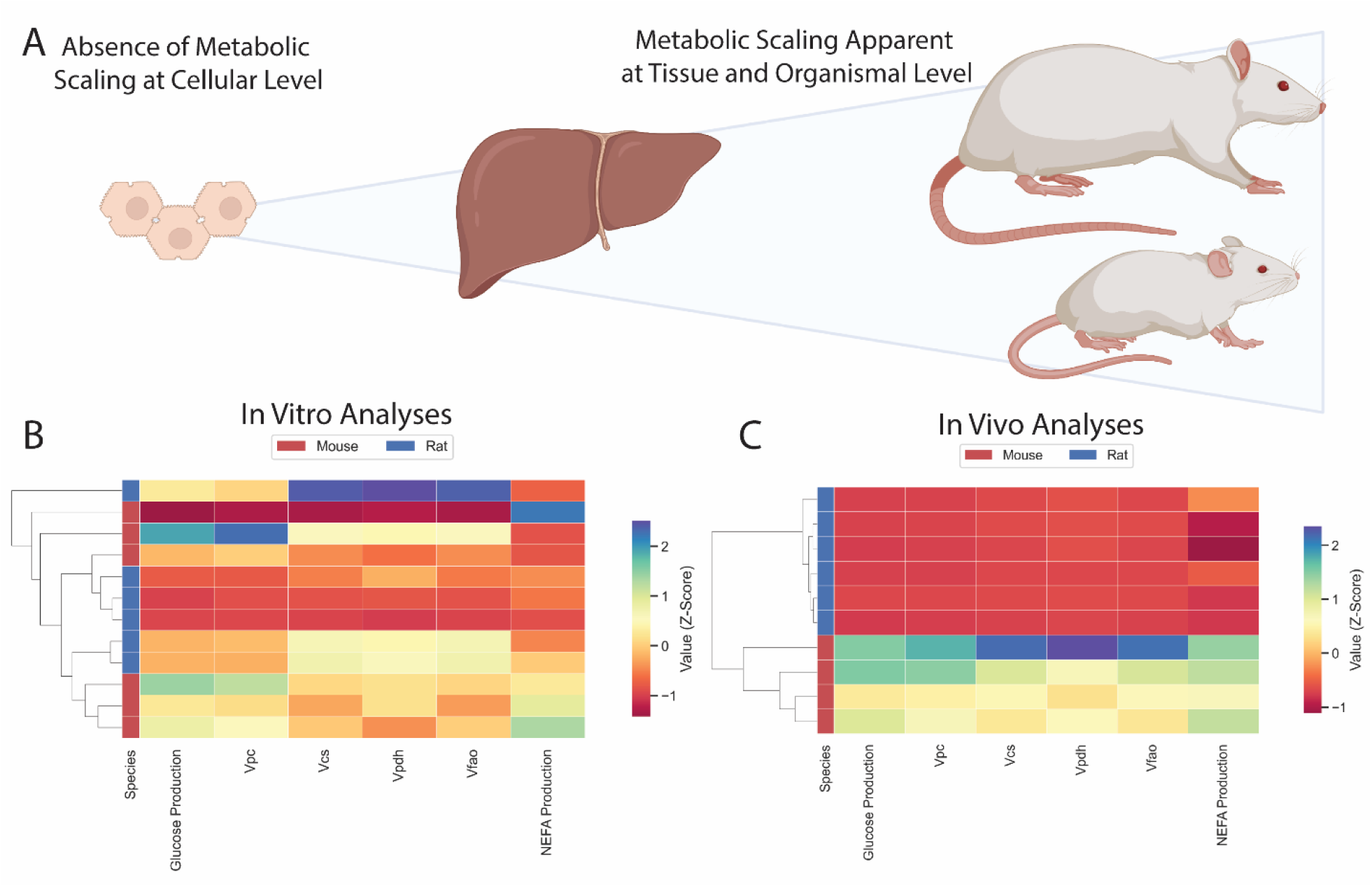
Comparison of *in vitro* and *in vivo* results. (A) Study workflow. (B) Clustering heatmap demonstrating the absence of metabolic scaling *in vitro*. (C) Clustering heatmap demonstrating metabolic scaling *in vivo*. In panels (B) and (C), mouse and rat color legends correspond to the species label attached to the dendrogram on the leftmost of each graph. Vpc = pyruvate carboxylase flux, Vcs = citrate synthase flux, Vpdh = pyruvate dehydrogenase flux, Vfao = fatty acid oxidation, NEFA = non-esterified fatty acid concentrations.

## Discussion

Oxygen consumption has been shown to scale inversely with body mass in species ranging in mass across 20 orders of magnitude, from 10^−14^ to 10^6^ grams (Ernest et al., 2003; Gillooly et al., 2001; Kleiber, 1932; Anastassia M Makarieva et al., 2005; Makarieva et al., 2008; Savage et al., 2004; West et al., 2002). This is perhaps best studied in mammals, but is a highly conserved phenomenon, having also been shown to occur in prokaryotes (Fenchel and Finlay, 1983; A. M. Makarieva et al., 2005; Anastassia M Makarieva et al., 2005; Makarieva et al., 2008; Moses et al., 2008), plants (A. M. Makarieva et al., 2005; Mori et al., 2010; Reich et al., 2006), insects (Chown et al., 2007; Maino and Kearney, 2014; A. M. Makarieva et al., 2005), fish (Clarke and Johnston, 1999; Gjoni et al., 2020; Rubalcaba et al., 2020), and birds (Glazier, 2008; Hudson et al., 2013; A. M. Makarieva et al., 2005). However, a major limitation of prior studies in this field has been that observations have been largely limited to oxygen consumption and caloric intake, leaving other metabolic processes unexplored. While oxygen consumption – reflecting mitochondrial oxidation of carbohydrates and/or fatty acids – is a crucial process for energy generation and reflects the steady-state basal metabolic rate, relying only on this variable, is a crude approach to examine metabolism and can fail to detect differences in substrate contributions to mitochondrial oxidation and cytosolic processes such as gluconeogenesis, in addition to the genes that set the capacity for these processes. This study sought to address this issue by examining the generalizability of metabolic scaling.

By combining mRNA expression measurement across a 30,000-fold range in body mass with isotopic tracer techniques to examine metabolic flux rates *in vivo, ex vivo*, and *in vitro* in mice and rats, this work provides new insights regarding inter- and intra-species metabolic scaling. Hepatic mitochondrial oxidation and the contributions of glucose and fatty acids to mitochondrial TCA cycle flux scaled similarly to cytosolic gluconeogenesis and white adipose tissue lipolysis *in vivo*, demonstrating systemic conservation of metabolic scaling. These data suggest that metabolic scaling may be subject to systemic signals to coordinate metabolic rates. Consistent with this interpretation is the fact that most metabolic processes tested – mitochondrial oxidation, the contributions of glucose and fatty acids to TCA cycle flux, glucose production, and lipolysis – scaled similarly, despite the different cellular compartments and tissues responsible for these fluxes.

Interestingly, the scaling of GPT and ADIPOR1 further suggest that there is dependence on extra-hepatic organs in the scaling of *in vivo* gluconeogenesis and fatty acid oxidation: that is, skeletal muscle supply of alanine for the liver mediated glucose-alanine cycle and adipose-tissue derived adiponectin signaling. These findings also suggests that the scaling of mitochondrial mass (Porter and Brand, 1995) or mitochondrial proton leak (Porter and Brand, 1993) cannot fully explain metabolic scaling.

The possibility that gene expression, as reflected by mRNA abundance, may also scale with body mass has not been previously addressed. We observed that expression of key genes in glycolysis, gluconeogenesis, fatty acid metabolism, NAD synthesis and transport, mitochondrial oxygen consumption, in addition to protection from oxidative damage, scales with body mass. More compelling, however, is the fact that those genes for which scaling of expression with body mass is observed, are not randomly distributed across the genome. Rather, the collection of genes that scale is enriched for genes related to metabolic processes, and whose proteins’ functions are constrained by supply of substrate, NAD, ATP, or oxygen. The notion that body mass is a variable related to expression level of certain genes has not previously been considered as an aspect of metabolic scaling, and when combined with isotope-based demonstration of flux through these pathways, yields new insight into the systems regulation of scaling.

It is also informative to consider the settings in which metabolic scaling was not observed (oxygen consumption, mitochondrial oxidation, lipolysis, and glucose production in hepatocytes). The differences between findings *in vitro* (where mouse and rat hepatocytes showed no significant difference in any cytosolic or mitochondrial flux) and *in vivo* observations (where livers exhibited robust metabolic scaling, with all fluxes in mice faster than those of rats) were striking, and emphasize the need to employ tracer methods *in vivo* to generate a comprehensive picture of the physiologic role of metabolic scaling.

Our findings emphasize that measurement of oxygen consumption *in vitro* may fail to detect any influence of scaling processes present *in vivo*. Glucose production was three-fold higher in mouse liver slices relative to rat liver slices, but did not significantly differ between plated hepatocytes from mice and rats, suggesting a role for extrinsic signals in metabolic scaling.

Taken together, the findings of this study show that the phenomenon of metabolic scaling extends to processes beyond oxygen consumption and caloric intake, and are bookended by scaling not only of gene expression, but also of a plethora of cytosolic and mitochondrial fluxes *in vivo*. The mechanisms underlying the phenomenon remain obscure, and deserve investigation as a fundamental question in biology (Hatton et al., 2019; Kolokotrones et al., 2010; White and Seymour, 2003).

### Ideas and Speculation

These data have implications for metabolic regulation during hibernation, in which metabolic rates are reproducibly and markedly but reversibly suppressed (Jansen et al., 2021, 2019; Tøien et al., 2011). Metabolic scaling phenomena may also have medical relevance. Recent reports provide data consistent with “Peto’s paradox” by showing that cancer is not more prevalent in larger, long-lived organisms than smaller ones (Anastassia M Makarieva et al., 2005; Vincze et al., 2022), despite the fact that more cells are at risk for transformation, and over longer periods of time. Metabolic scaling provides a potential explanation: the slower rates of metabolic processes (including oxygen consumption) and reduced oxidative damage in larger animals imply lower rates of proliferation and carcinogenic DNA damage, so that while more cells are at risk in large animals, carcinogenesis proceeds more rapidly in smaller ones.

## Materials and Methods

### Rodents

Male C57bl/6J mice and Sprague-Dawley rats were obtained at 8 weeks of age from Jackson Laboratories and Charles River Laboratories, respectively, and given *ad lib* access to regular chow and water. Mice underwent surgery under isoflurane anesthesia to place catheters in the jugular vein (mice) and in both the jugular vein and carotid artery, with the tip of the arterial catheter advanced into the right atrium of the heart. After a week of recovery and confirmation that the animals had regained their pre-surgical body weight and following a 24 hour fast, rodents underwent the *in vivo* tracer studies described below. Rodents used for hepatocyte studies were fed *ad lib* until isoflurane euthanasia and liver isolation as described in the *in vitro* studies below.

### Gene Expression Analysis

All liver raw transcriptomics data were obtained from Array Express (Brazma et al., 2003), and were all preprocessed using the same methods. Cattle, monkey, and rat were obtained from E-MTAB-4550, mouse from E-MTAB-5166, and human from E-MTAB- 6814. Two replicates from each species were used. Raw counts from each species were normalized to counts per million (CPM) and were then TMM-normalized to account for differences in sequencing depth as well as transcript length across species and scanners using the R package edgeR (Robinson et al., 2010). All transcript homologues were converted to human gene names using ENSEMBL in the biomaRt R package (Durinck et al., 2009). For the analyses in Figure 5, genes were filtered to those that followed the allometric scaling pattern mouse > rat > monkey > human > cattle. This gene list was put into EnrichR and KEGG pathway enrichment analysis was performed (Kuleshov et al., 2016). Genes in this list that followed the scaling pattern were then filtered based on whether they met one of six Reactome metabolic superpathways related to metabolism of amino acids, carbohydrates, energy, lipids, tricarboxylic acid cycle, and vitamin cofactors (Jassal et al., 2020). A clustering heatmap was performed on these genes using the Seaborn Python package (Waskom, 2021). Genes were plotted again upon this map using the Seaborn Python package. To assess for differences in gene expression in genes displayed on the metabolic map, a one-way ANOVA with a Bonferoni correction for multiple comparisons (thus an adjusted p-value threshold for this test was 0.01 from 0.05 for significance) using the Python Scipy package (Virtanen et al., 2020).

### *In Vitro* Tracer Analysis

Primary hepatocytes were isolated by the Yale Liver Center’s Cell Isolation Core, plated in a six well plate (4.0×10^5^ cells per well), and allowed to recover for six hours at 37°C in substrate replete media (DMEM high glucose containing 10% FBS, 2% penicillin–streptomycin, 100 nM dexamethasone, 10 mM HEPES, and 1 nM insulin). The attached cells were then washed once in PBS and incubated overnight in low-glucose culture media (DMEM low glucose containing 10% FBS, 2% penicillin–streptomycin and 10 mM HEPES) for glucose production assays, or serum-free low-glucose culture medium (DMEM low glucose supplemented with 0.5% fatty acid free BSA, 2% penicillin–streptomycin and 10 mM HEPES) for lipolysis assays. Following the overnight incubation, for glucose production assays, cells were washed twice with PBS and the media was replaced with 2 ml of substrate replete glucose production medium (DMEM no-glucose base media containing 0.5% fatty acid free BSA, 20 mM sodium lactate (50% 3-^13^C), 2 mM sodium pyruvate, 2 mM GlutaMAX, MEM non- essential amino acids, and 10 mM HEPES). After a six hour incubation at 37°C, the media was collected. Glucose concentrations were measured as described in the Biochemical Analysis section, and normalized to total protein measured using the bicinchoninic acid (BCA) assay. For lipolysis assays, cells were washed twice with PBS and the media was replaced with a fresh 2 ml volume of low-glucose culture medium. Cells were incubated at 37°C for six hours, after which the media was collected and non-esterified fatty acid concentrations measured as described in the Biochemical Analysis section. The lipolysis rate was calculated after normalizing to total protein concentrations determined using the BCA assay. Flux ratios were measured using Equations 5-12, and back-calculated from net glucose production determined by measuring the glucose concentration in the media using the Sekisui Glucose Assay and assuming a linear rate of net glucose production during the 6 hour incubation.

### Glucose production in liver slices

Mice and rats were fasted overnight (16 hr) and sacrificed. Rodent livers were extracted and washed in Krebs-Henseleit buffer (KHB) containing 550 mM sodium chloride, 23 mM potassium chloride, 6.3 mM calcium dichloride, 10 mM magnesium sulfate, and 6.9 mM sodium phosphate monobasic. Rodent livers were cored into smaller bits using the Alabama R&D tissue coring press. Cored livers were sliced to the thickness of 230 micron, the lowest setting on the Alabama R&D tissue slicer. Liver slices were transferred to 24 well plates containing gluconeogenesis media (DMEM without glucose but with 20 mM lactate, 2 mM pyruvate, 10 mM HEPES, and 44 mM sodium bicarbonate) (400 ul for mouse livers and 500 ul for rat livers). The 24 well plates were placed in a tissue culture incubator (5% CO_2_) and shaken at 80 RPM for 6 hours. At the end of the 6 hr incubation, liver slices and media were collected. Liver slice weights were measured on a scale. Glucose concentrations from liver slice media were measured using the Sekisui Glucose Assay. Glucose concentrations was normalized to liver weights measured in milligrams and microliters of gluconeogenesis media.

### *In vitro* oxygen consumption analysis

Primary hepatocytes were isolated from *ad lib* fed mice and rats by the Yale Liver Center’s Cell Isolation core and plated recovery media as described previously(Camporez et al., 2013) in 24-well XF24 V7 cell culture plates coated with type I collagen. After 6-8 hours of recovery at 37°C in 5% CO_2_, cells were washed twice with PBS and the media was replaced with low-glucose culture media (DMEM base medium containing 5 mM glucose, 2 mM glutamine, and non-essential amino acids, pH 7.4), in which cells were cultured overnight. The next morning, as we have previously described(Perry et al., 2020), cells were washed twice with PBS and the media was replaced with 500 µL XF24 assay medium (DMEM base medium containing 5.5 mM glucose, 1 mM pyruvate, and 2 mM glutamine, pH 7.4) and equilibrated at 37°C for 60 min. The Seahorse XFe 24 Analyzer was used to perform a mitochondrial stress test: after three baseline measurements of O_2_ consumption (10 min apart), oligomycin (an inhibitor of ATP synthase) was injected, and three subsequent measures of O_2_ consumption were performed using a 4 min mix/2 min wait/4 min measure protocol. Next, the uncoupler 2-[2-[4-(trifluoromethoxy)phenyl]hydrazinylidene]-propanedinitrile (FCCP) was injected to dissipate the proton gradient, with three O_2_ consumption measurements taken as described above. Finally, rotenone (0.5 µM) and antimycin (10 µM) were injected to inhibit Complexes I and III, respectively. Oxygen consumption was normalized to total protein measured using the Pierce BCA Protein Assay.

### *In Vivo* Tracer Analysis

Mice and rats received a 3X (5 min) primed-continuous infusion of [3- ^13^C] sodium lactate (4.5 mg/kg body weight/min), [1,2,3,4,5,6,6-^2^H_7_] glucose (1.0 mg/kg/min), and [U-^13^C_16_] potassium palmitate (0.8 mg/kg/min) for 120 min. Tracers were infused into the jugular vein (mice) or the right atrium (rats) to ensure systemic delivery. After 120 min, blood was collected from the tail vein (mice) or from the jugular venous catheter (rats) and centrifuged to obtain plasma, then animals were sacrificed with IV pentobarbital and their livers freeze-clamped in liquid nitrogen.

### Biochemical analysis

In plated hepatocyte and liver slice studies, glucose concentrations in the cell media were determined using the Sekisui Glucose Assay. Non-esterified fatty acid concentrations in the hepatocyte media were measured using the Sekisui Non-Esterified Fatty Acid kit.

Glucose production in hepatocytes was determined by measuring the glucose concentration in media after 6 hr of incubation. Glucose enrichment in hepatocytes, plasma, and livers was measured by gas chromatography/mass spectrometry (GC/MS). Samples were deprotonized using 1:1 barium hydroxide:zinc sulfate (100 µl for cells and 300 µl for ∼100 mg liver samples, which were subsequently homogenized using a TissueLyser) and derivatized with 50 µl 1:1 acetic anhydride:pyridine. After 20 min heating to 65°C, 50 µl methanol was added, and glucose enrichment ([^13^C_1_], [^13^C_2_] and, in plasma, [^2^H_7_] (all using the Chemical Ionization mode), [4,5,6- ^13^C_1_] and [4,5,6-^13^C_2_] (both using the electron ionization mode)) were measured by GC/MS.

[^13^C] alanine enrichment was measured by GC/MS (Perry et al., 2016). We have previously shown that flux through pyruvate kinase and malic enzyme – which would add [^13^C] label to carbon 2 of alanine – is minimal under fasting conditions in normal rats (Perry et al., 2016); therefore, the measured [^13^C] alanine enrichment can be attributed entirely to [3-^13^C] lactate enrichment. [^13^C] malate enrichment was also measured by GC/MS (Perry et al., 2017b). Total malate enrichment and C1C2C3 malate enrichment were measured and the C2 + C3 malate enrichment was determined according to **Table S1**, equations 1 and 2, which relies on the assumption that the C4 enrichment is approximately equal to the C1 enrichment of malate (Perry et al., 2017b). Total and C4C5 glutamate enrichment was measured by LC-MS/MS (Perry et al., 2018). With ^13^C lactate infusion, no label enters glutamate carbon 5, so all label in the C4C5 fragment was assumed to be label in glutamate C4.

### *In vivo* flux analysis

Endogenous palmitate and glucose production were determined using equation 3 in **Table S1** (Perry et al., 2017a) comparing plasma enrichment to the infused tracer enrichment, measured using gas chromatography/mass spectrometry (GC/MS) as described in the Biochemical Analysis section. In the fasted/substrate depleted and therefore glycogen-depleted state (Perry et al., 2018), endogenous glucose production can be attributed entirely to gluconeogenesis (equation 4). Based on equations previously described (Perry et al., 2017b), and after verifying minimal renal and hepatic bicarbonate enrichment by GC/MS (Perry et al., 2020, 2017b), we measured the whole-body ratio of phosphoenolpyruvate carboxykinase (PEPCK) flux (i.e. gluconeogenesis from pyruvate) to total gluconeogenesis by mass isotopomer distribution analysis (equations 5 and 6), correcting for any [^13^C_2_] glucose synthesized from [^13^C_2_] trioses (equation 7). At steady state, V_PEPCK_ is equal to the sum of pyruvate kinase flux and pyruvate carboxylase flux (V_PK_ + V_PC_); based on our previously published data, under fasting conditions pyruvate kinase flux is minimal, less than 10% of pyruvate carboxylase flux (Perry et al., 2016). Therefore, we can assume that the rate of gluconeogenesis from pyruvate (i.e. V_PEPCK_) is approximately equal to V_PC_.

Next, we measured the ratio of pyruvate carboxylase anaplerosis to citrate synthase flux (V_PC_/V_CS_) using the enrichment of liver alanine, malate, and glutamate (equation 8). This equation, in which pyruvate cycling is again assumed minimal, is derived in detail in our recent publication (Perry et al., 2017b).

We measured the fractional contribution of glycolytic carbons to the TCA cycle (i.e. pyruvate dehydrogenase flux relative to citrate synthase flux, V_PDH_/V_CS_) (equation 9) using the model and assumptions we (Befroy et al., 2014; Perry et al., 2018, 2016; Song et al., 2020) and others (Alves et al., 2011; Petersen et al., 2016, 2015) have described. Finally, absolute turnover rates (equations 10-13) were determined, utilizing the ratios measured with equations 5, 8, and 9, and the absolute gluconeogenesis rate measured using equations 3 and 4.

### Statistical considerations, power calculations, and statistical analysis

The sample size (n=4-6 *in vitro* replicates or animals *in vivo*) was calculated to supply 80% power at α=0.05 to detect the expected two-fold difference with 50% variance. Power calculations were performed using the ClinCalc online calculator. The *in vivo* studies compared 4-6 biological replicates (unique animals), the liver slice studies compared 2-4 biological replicates, and the *in vitro* studies compared 2 biological replicates, with 3 technical replicates (separate wells) from each. Rodents were excluded from analysis following tracer studies if they failed to respond immediately (i.e. immediate euthanasia, within 3 seconds) to IV pentobarbital. No samples were excluded from analysis in the *in vitro* or *ex vivo* studies. Randomization was not possible during the *in vivo* studies because of the readily apparent differences between mice and rats, but all analyses were performed by investigators who were blinded as to species.

## Acknowledgments

The authors thank Traci LaMoia for her assistance in performing the liver slice experiments, Kathy Harry of the Yale Liver Center for isolating hepatocytes, and members of the Perry and Pollak labs for helpful discussions. This study was funded by grants from the U.S. Public Health Service (K99/R00 R00CA215315 and R37 258262A1-01 [to R.J.P.], T32-GM0007324 [supporting N.D.A.], T32GM136651 [supporting B.P.L.], and P30DK034989 [supporting the Yale Liver Center]).

## Supplementary Figures and Table

**Figure S1.**
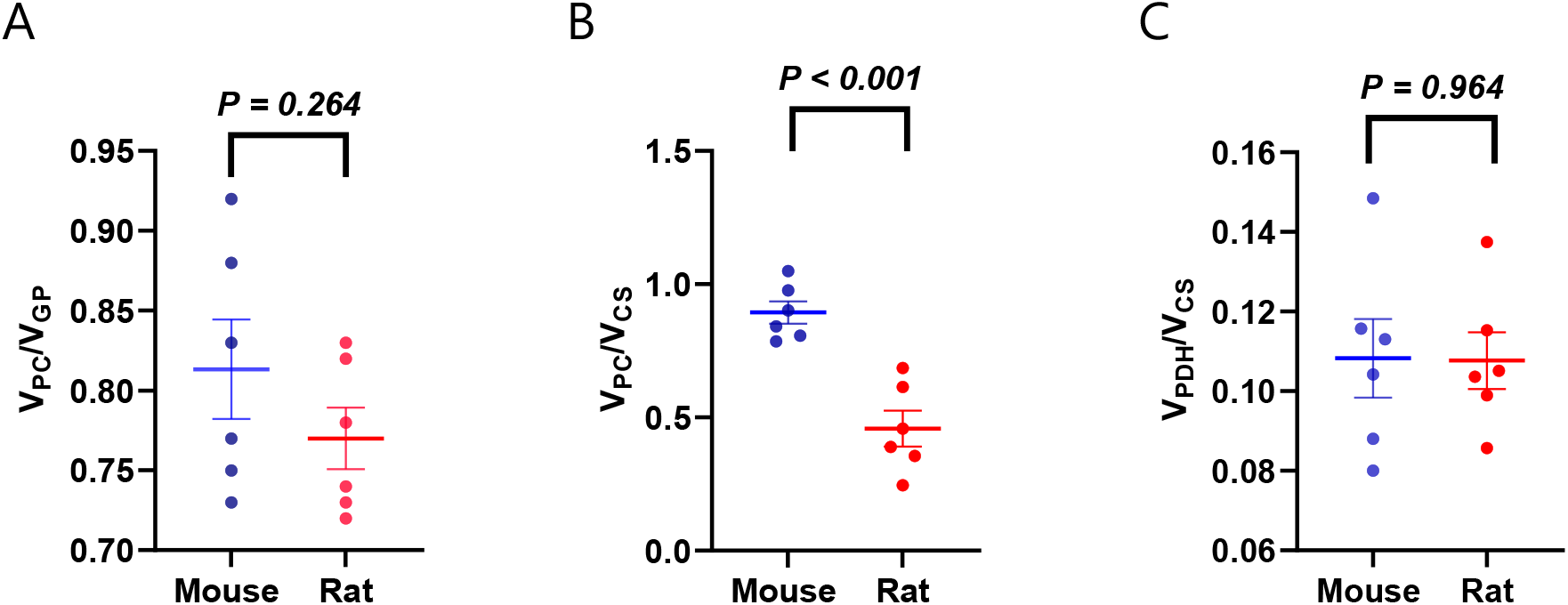
Flux ratios in plated hepatocytes. (A) V_PC_/V_GP_. (B) V_PC_/V_CS_. (C) V_PDH_/V_CS_.

**Figure S2.**
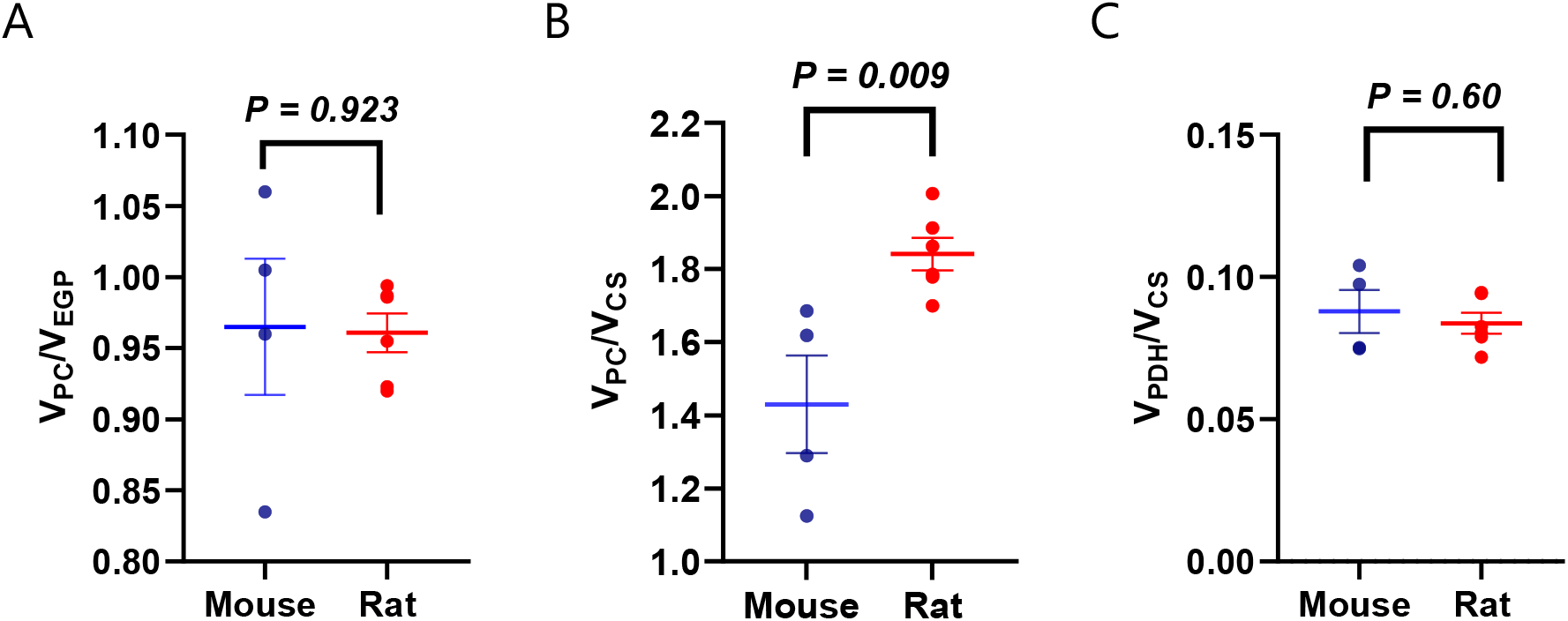
Flux ratios *in vivo*. (A) V_PC_/V_EGP_. (B) V_PC_/V_CS_. (C) V_PDH_/V_CS_.

**Table S1.** Flux ratios and absolute rates. measured in mice infused with [3-^13^C] lactate. APE indicates the atom percent enrichment (in animals infused with ^13^C tracer), TCA denotes the tricarboxylic acid cycle, and GNG denotes gluconeogenesis. By convention, V_a_ represents the flux through pathway *a*.

| Equation | Interpretation |
| --- | --- |
| 1. $^{13}\text{C4 malate} = \text{Total } ^{13}\text{C malate} - ^{13}\text{C1C2C3 malate}$ | [4-$^{13}\text{C}$] malate, equivalent to [1-$^{13}\text{C}$] malate |
| 2. $^{13}\text{C2C3 malate} = \text{Total } ^{13}\text{C malate} - 2 * (^{13}\text{C4 malate})$ | Enrichment in carbons 2 and 3 of malate |
| 3. $\text{Turnover} = \left( \frac{\text{Tracer APE}}{\text{Plasma APE}} - 1 \right) * \text{Infusion rate}$ | Whole-body endogenous glucose or palmitate production |
| 4. $\text{GNG} = \text{Glucose turnover}$ | Whole-body gluconeogenesis |
| 5. $\frac{V_{\text{PEPCK}}}{V_{\text{GNG}}} \sim \frac{V_{\text{PC}}}{V_{\text{GNG}}} = \frac{[^{13}\text{C}_2]\text{glucose}}{\text{XFE}^2}$ | Fraction of gluconeogenesis derived from pyruvate |
| 6. $\text{XFE} = \frac{1}{1 + \frac{[^{13}\text{C}_1]\text{glucose}}{2 * [^{13}\text{C}_2]\text{glucose}}}$ | Fractional triose enrichment |
| 7. $\text{Corrected } [^{13}\text{C}_2]\text{glucose} = \text{Measured } [^{13}\text{C}_2]\text{glucose} - 2 * [\text{C4C5C6} - ^{13}\text{C}_2]\text{glucose}$ | Doubly-labeled glucose arising from the condensation of two singly labeled trioses, correcting for doubly labeled glucose arising from one doubly labeled triose condensing with an unlabeled triose |
| 8. $\frac{V_{\text{PC}}}{V_{\text{CS}}} = \frac{[5-^{13}\text{C}]\text{glucose}}{2 * [4-^{13}\text{C}]\text{glucose}}$ | Rate of pyruvate carboxylase anaplerosis relative to TCA cycle flux |
| 9. $\frac{V_{\text{PDH}}}{V_{\text{CS}}} = \frac{[4-^{13}\text{C}]\text{glutamate}}{[^{13}\text{C}]\text{alanine}}$ | Fractional contribution of glucose to the TCA cycle |
| 10. $V_{\text{PC}} = \frac{V_{\text{PC}}}{V_{\text{GNG}}} * V_{\text{GNG}}$ | Absolute rate of gluconeogenesis from pyruvate |
| 11. $V_{\text{CS}} = \left( \frac{V_{\text{PC}}}{V_{\text{CS}}} \right)^{-1} * V_{\text{PC}}$ | Absolute TCA cycle flux |
| 12. $V_{\text{PDH}} = \frac{V_{\text{PDH}}}{V_{\text{CS}}} * V_{\text{CS}}$ | Absolute rate of glycolytic carbon entry into the TCA cycle |
| 13. $V_{\text{FAO}} = V_{\text{CS}} - V_{\text{PDH}}$ | Absolute rate of entry of carbons from oxidized fatty acids into the TCA cycle |

